# DNMT3b Dysfunction Promotes DNA cleavages at Centromeric R-loops to Increase Centromere Instability

**DOI:** 10.1101/2020.06.04.133272

**Authors:** Hsueh-Tzu Shih, Wei-Yi Chen, Hsin-Yen Wang, Hsien-Da Huang, Chih-Hung Chou, Zee-Fen Chang

## Abstract

This study investigates how DNA methyltransferase 3b (DNMT3b) dysfunction causes genome instability. We showed that in DNMT3b deficient cells, R-loops contribute to prominent γH2AX signal, which was mapped to repetitive satellite sequences including centromere regions. By ChIP and DRIP analyses, our data revealed that centromeric R-loops in DNMT3b deficient cells are removed by XPG/XPF, thus generating DNA breaks in centromeres to increase mitotic aberration. In immunodeficiency-centromeric instability-facial anomalies (ICF) patient cells carrying the loss-of-function mutation at *DNMT3b*, knockdown of XPG/XPF in ICF cells also reduces DNA breaks in centromere while bringing up centromeric R-loop to the level similar to that in wild-type cells. These results suggest that DNMT3b has a critical function in preventing XPG/XPF-mediated cleavages at centromeric R-loop sites. Finally, we showed the involvement of non-homologous end-joining repair at centromeric sites in ICF cells. Thus, DNA cleavages at centromeric R-loops with error-prone repair undermine centromere stability in ICF cells.

## INTRODUCTION

DNA methyltransferase DNMT3b carries out *de novo* DNA methylation and is essential for mammalian development (1,2). The murine embryonic fibroblasts derived from *DNMT3b* knockout embryo display DNA damage and chromosome instability (1,3), suggesting the critical function of DNMT3b in genome stability. Loss-of-function mutation in *DNMT3b* is specifically found in a rare human genetic disorder, immunodeficiency-centromeric instability-facial anomalies (type 1 ICF) syndrome (4-6). It has been shown that DNMT3b is recruited to GC-rich (peri-)centromere regions by interacting with centromere protein CENP-C for maintaining chromosome stability (7). In accordance, the major genome regions affected by the loss of DNMT3b function in ICF are non-coding repetitive elements surrounding centromeres, where GC regions are hypomethylated (8), coinciding with centromeric DNA breaks observed in ICF cells (9,10). However, it is still an open question how DNMT3b dysfunction increases DNA damage and chromosome instability.

R-loop is a three-stranded nucleic acid structure consisting of an RNA-template DNA hybrid and a non-template single-stranded DNA (11). In mitosis, centromeric domain remains transcriptionally active (12). It has been reported that centromeric R-loop has a function in mediating ATR binding for checkpoint control in mitosis (13), suggesting the beneficial role of R-loops in a cell. However, R-loops are also considered as the source of DNA damage (14,15). R-loops can be resolved by helicases, translocase, or removed by RNase H (16-18). Without resolution, R-loops lead to DNA double-strand breaks (DSBs) through replication fork collapse (14) or DNA cleavage by endonucleases XPG and XPF (19,20). In addition, it has been shown that chromatin modifications regulate R-loop-induced genome instability (21-24). Since DNMT3b is an epigenetic factor that regulates transcription, we asked the question whether the formation or processing of R-loops is involved in DNMT3b dysfunction-mediated DNA damage and centromere instability.

In this study, we address the question by using human colon cancer HCT116 BKO cells in which the allele of DNMT3b is disrupted by homologous recombination, which results in less than 3% reduction in genome methylation (25), and ICF cells that carry loss-of-function mutation of *DNMT3b*. Our data suggest that *DNMT3b* knockout in HCT116 or the loss-of-function mutation of DNMT3b in ICF lymphocytes increases R-loops processing by XPG/XPF-mediated cleavages, thereby generating more DNA breaks in centromere. Finally, we examined the choice of DNA double-strand break (DSB) repair pathway in centromeric breaks in ICF cells. Our data indicate that DSB repair at centromere sites in ICF cells is mainly through non-homologous end-join process (NHEJ), which is an error-prone repair (26). Here, we proposed that DNMT3b dysfunction promotes XPG/XPF-mediated DNA breaks at centromeric R-loops sites, where mutagenic repair of DSBs via NHEJ increases centromeric shortening and fusion that undermine centromere stability in ICF cells.

## RESULTS

### DNMT3b deficiency increases R-loops-dependent DNA damage signals by XPG/XPF in HCT116 cells

To access the importance of DNMT3b in genome instability, the levels of DSBs in HCT116 and BKO cells were compared by γH2AX IF staining. Data showed that γH2AX signal was very prominent all over the nuclei in BKO cells (Fig. 1a), so as other DNA damage response signals, including phospho-ATM, -Chk2 and -Chk1 (Fig. 1b). After enforced expression of HA-RNase H1, γH2AX signal in BKO cells diminished (Fig. 1c). Since R-loops are the results of transcription, we then treated BKO cells with α-Amanitin (20 μg/ml) or cordycepin (50 μM) for 6 h, and found that γH2AX intensity was clearly reduced (supplementary Fig.S1). These results suggest that R-loops contribute to the increase of DNA damage in BKO cells. A previous report has shown that transcription-coupled nucleotide excision repair (TC-NER)-mediated endonuclease cleavage of unprocessed R-loops promotes genome instability (19,20). We then depleted Cockayne Syndrome B (CSB) protein, a critical TC-NER factor in BKO cells. The overall γH2AX intensity was reduced in BKO cells after knockdown of CSB, suggesting the involvement of NER in making DSBs in these cells (Fig. 1d). Since XPF and XPG are the endonucleases of NER, we further depleted them in BKO cells. Knockdown of XPG or XPF alone had no significant effect on the level of γH2AX staining (supplementary Fig.S2). However, XPF/XPG-double depletion abolished γH2AX in BKO cells (Fig. 1e). This implies that XPG or XPF cleavage is still sufficient to cause DSBs. Presumably, knockdown of either XPG or XPF should be sufficient to prevent R-loop-mediated DSBs. Therefore, it is possible that XPG or XPF-mediated nicks are turned to DSBs through DNA replication (27). By DNA fiber analysis, BKO cells did show a marked increase in replication stress (Fig. 1f). Here, we conclude that DNMT3b deficiency promotes DNA damage due to NER-mediated R-loops processing.

**Figure 1.**
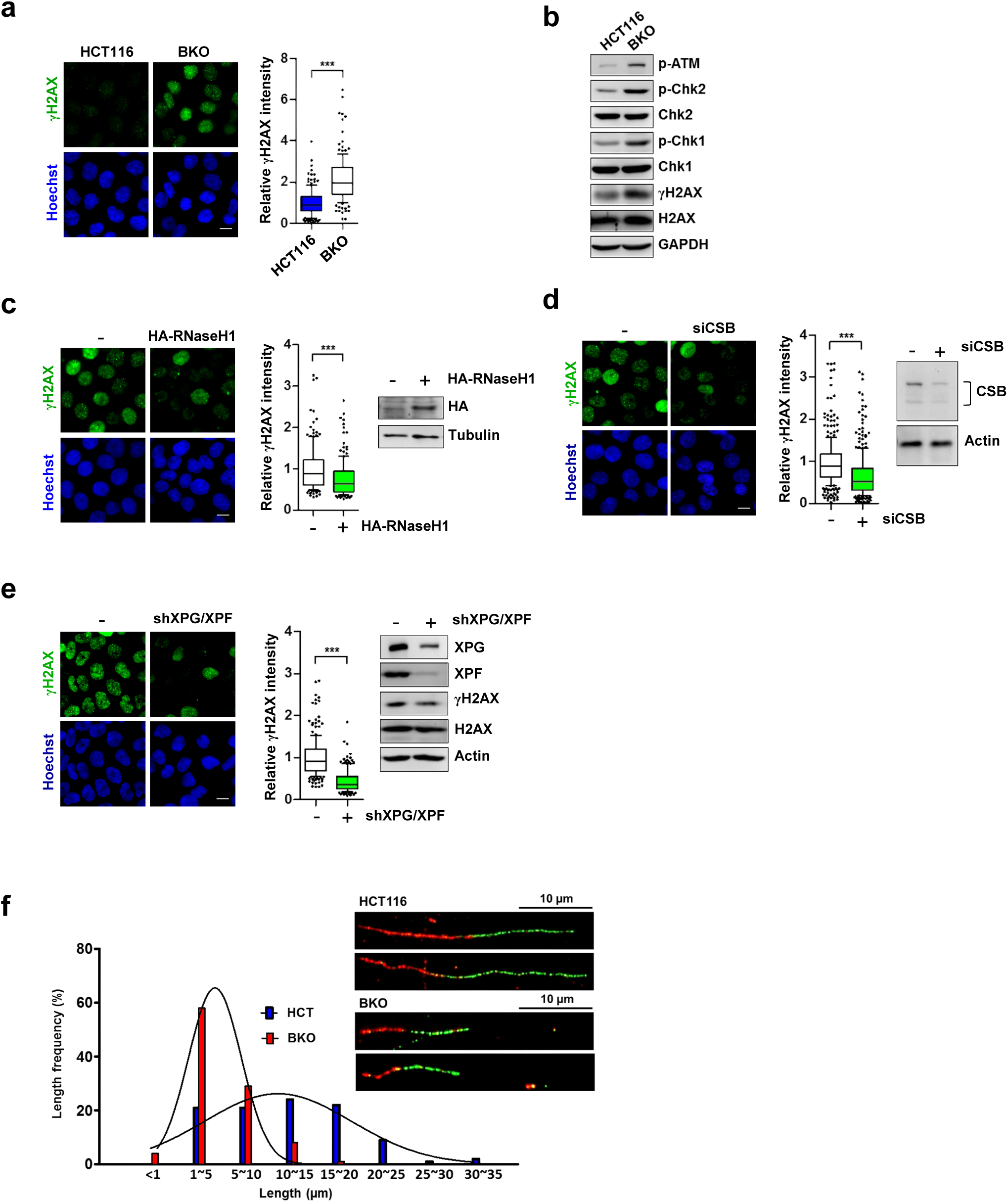
DNA damages are generated by XPG/XPF-mediated R-loops removal in DNMT3b deficient HCT116 cells. (a-b) HCT116 and BKO cells were fixed for (a) γH2AX IF staining (scale bar, 10 μm). Fluorescent intensity of γH2AX in cells (N>150) was quantitated by Image J from three independent experiments, and relative intensity is expressed, *\*\*\**P< 0.001 by Mann-Whitney test. (b) Western blot analysis of DNA damage response markers. (c) γH2AX IF staining in BKO cells that were infected with retrovirus of empty vector or HA-RNase H1. Fluorescent intensity of γH2AX was quantitated in cells (N>150) from three independent experiments and expressed as described in (a). The expression of HA-RNase H1 was indicated by western blot. (d) BKO cells were transfected by siCSB for 3 days for γH2AX IF staining (scale bar, 10 μm). Data from three independent experiments are expressed as described in (a). The knockdown of CSB was indicated by western blot. (e) BKO cells were infected with shXPG virus. After recovery and selection, cells were subsequently infected by virus of shXPF. Cells were recovered for 2 days prior to analysis. IF staining of γH2AX in cells (scale bar, 10 μm) and fluorescent intensity was quantitated (N>150) from three independent experiments. Relative intensity is expressed, *\*\*\**P< 0.001 by Mann-Whitney test. Western blot analysis of XPG, XPF, γH2AX, H2AX, and actin is shown in the right panel. (f) DNA fiber analysis of DNA replication track. HCT116 and BKO cells were pulse-labeled with CldU (red) followed by IdU (green) for DNA fiber analysis as described in materials and methods. Representative DNA fiber images are given. The length of IdU tract connected to CIdU labeled fiber was measured (N>100).

### XPG/XPF-mediated DSBs are mapped to repetitive satellite sequences

Since XPG/XPF knockdown markedly reduced γH2AX IF staining, we then wanted to identify R-loops-mediated DNA damage sites. HCT116 and BKO cells were used to perform ChIP-sequencing analysis using γH2AX antibody. This genome-wide search mapped DSBs sites at repetitive satellite sequences, including centromere and telomere, and ribosomal DNA (rDNA) in BKO cells (Fig. 2a). These data are correlated with transcription abnormalities previously observed at centromere in *DNMT3b*-depleted HCT116 cells (7), telomeres in ICF cells (28), and rDNA in *DNMT1/DNMT3b*-disrupted HCT116 cells (29). ChIP-qPCR confirmed higher levels of γH2AX binding at sequences of chromosome 1 centromere, telomere, and rDNA repeats in BKO than HCT116 cells. Importantly, the association of γH2AX with centromere, telomere, and rDNA repetitive sequences was reduced by XPG/XPF knockdown specifically in BKO cells (Fig. 2b). As a comparison, there were little differences in γH2AX binding at p21(*CDKN1A*) promoter in response to XPG/XPF knockdown in HCT116 and BKO cells. Therefore, XPG/XPF-mediated DNA breaks at repetitive satellite and rDNA sequences make a major contribution to the prominent DNA damage signals seen in BKO cells

**Figure 2.**
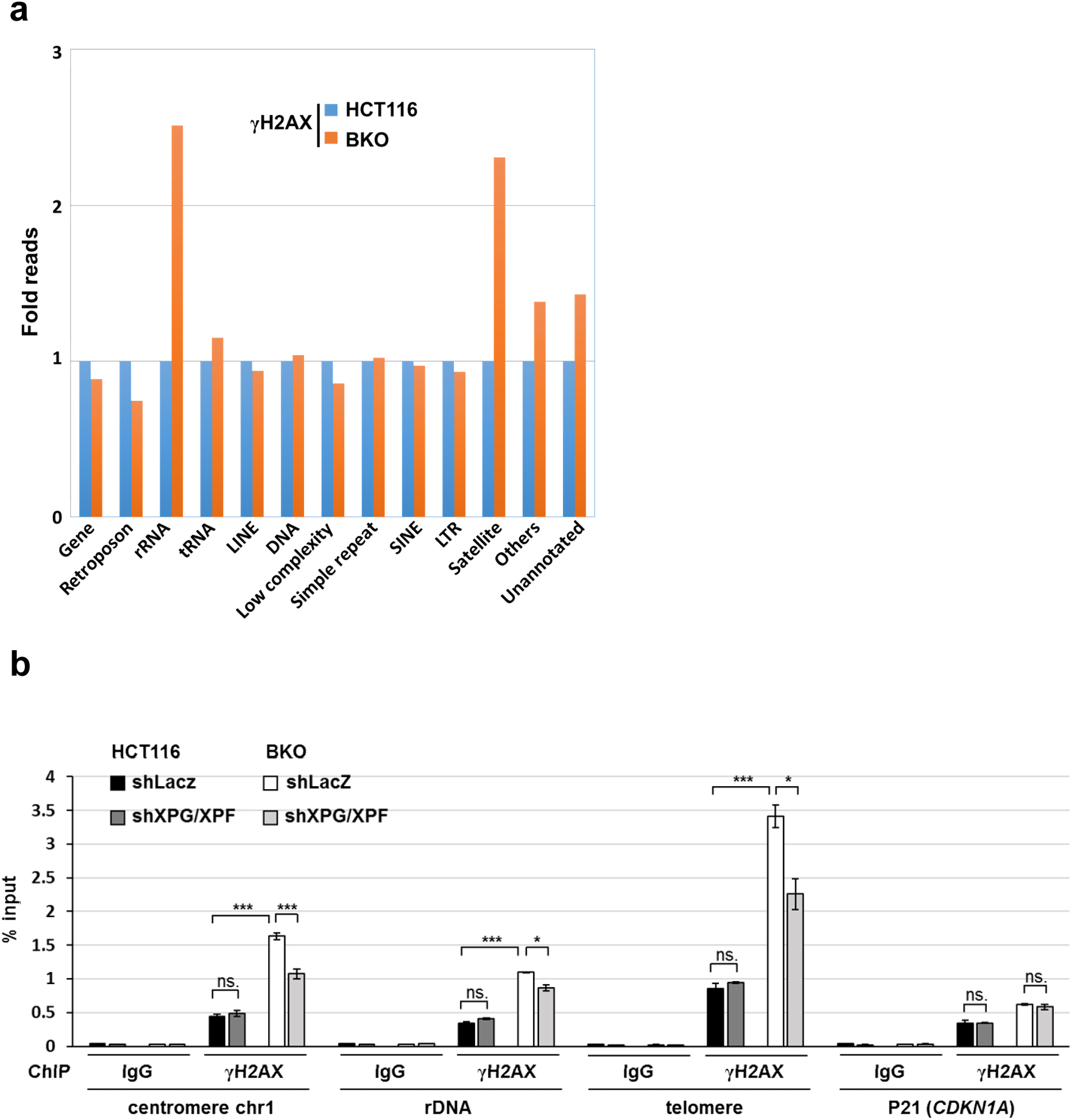
DNMT3b deficiency increases DSBs at centromere, telomere and rDNA sequences by XPG/XPF-mediated cleavages. (a) HCT116 and BKO cells were used for ChIP by γH2AX antibody, followed by genome-wide sequencing analysis. Plot shows the relative ChIP-seq reads of γH2AX at indicated genomic regions in BKO versus control HCT116 cells. Data were from two independent experiments. (b) HCT116 and BKO cells with or without XPG/XPF knockdown were used for γH2AX ChIP-qPCR analysis. Data are shown as percentage of input DNA in γH2AX antibody versus IgG control at centromere sequence of chromosome 1, rDNA, telomere sequence of chromosome 10 and p21 (*CDKN1A*) regions (mean ± SEM, n=3, *, ***P< 0.05, 0.001, ns: no significant by two-tailed unpaired Student’s t-test).

### The removal of centromeric R-loops by XPG/XPF processing contributes to mitotic defect in BKO cells

To know whether the level of centromeric R-loop is different in HCT116 and BKO cells and the changes in response to XPG/XPF knockdown, we performed DRIP analysis using DNA-RNA hybrid recognition antibody S9.6. All the samples were pretreated with and without RNase H, which removes all RNA transcripts, before immunoprecipitation by S9.6 antibody. DRIP data sensitive to RNase H pretreatment were considered to indicate the level of R-loop. The analyses revealed that the steady-state level of centromeric R-loop in HCT116 was indeed higher than that in BKO cells. XPG/XPF knockdown had no effect on increasing centromeric R-loop level in HCT116. As a contrast in BKO cells, XPG/XPF depletion brought up the level of centromeric R-loop higher than that in HCT116 cells (Fig. 3a).

**Figure 3.**
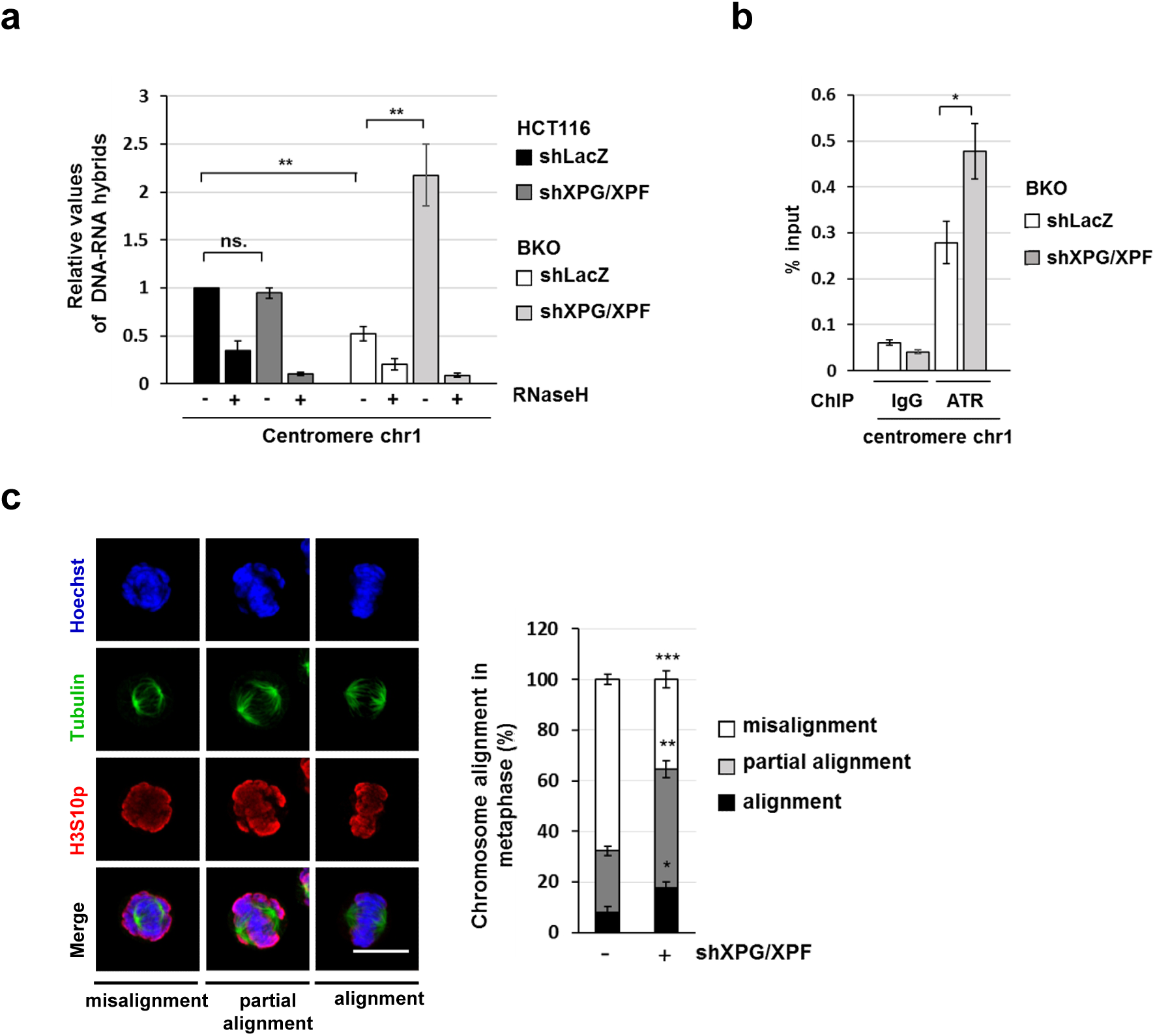
Mitotic defects in BKO cells were significantly reduced by XPG/XPF knockdown. (a) DRIP analysis in HCT116 and BKO cells with or without XPG/XPF knockdown. DNA samples were untreated (-) or treated (+) with RNaseH as indicated before immunoprecipitated by S.9.6 antibody. qPCR values of DNA-RNA hybrids at centromere sequence of chromosome 1 are normalized by IgG control. Data are expressed relative to that in HCT116 cells and presented as mean ± SEM of three independent experiments (**P< 0.01, ns: no significant by two-tailed unpaired Student’s t-test). (b) ATR ChIP analysis in mitotic arrested BKO cells with or without XPG/XPF knockdown. Data are shown as percentage of input DNA at centromere sequence of chromosome 1of ATR antibody versus IgG control (mean ± SEM, n=3, *P< 0.05, ns: no significant by two-tailed unpaired Student’s t-test). (c) Mitotic progression in BKO cells with or without XPG/XPF knockdown. Metaphase cells were defined by H3S10P and β-tubulin staining. Representative images of cell displaying aligned, misaligned, and partial misaligned metaphase chromatin are shown (left). More than 100 cells were counted in each experiment (means ± SEM, three independent experiments. *, **, *** P< 0.05, 0.01, 0.001, ns: no significant by two-tailed unpaired Student’s t-test).

It has been shown that ATR binding to centromeric R-loop serves as a mitotic checkpoint control (13). ATR-ChIP analysis revealed that XPG/XPF depletion did increase ATR binding to centromere in BKO cells (Figure 3b). Centromere integrity is important in chromosome segregation. We compared the chromatin alignment in metaphase in HCT116 and BKO cells. To this end, cells were treated with nocodazole overnight and washout for mitotic progression. Chromatin alignment in metaphase was evaluated by H3ser10 phosphorylation in cells containing polar spindles marked by tubulin staining. About 70% of mitotic BKO cells exhibited obvious aberrant chromatin alignment on the metaphase plate, while HCT116 cells had much less defects in chromatin alignment (supplementary Fig.S3). We further tested whether XPG/XPF knockdown affects mitotic progression in BKO cells. The results showed that the fraction of cells containing misaligned chromatin was significantly reduced by XPG/XPF knockdown (Fig. 3c), suggesting that XPG/XPF-mediated DNA damage contributes to mitotic aberration in BKO cells.

### Recapitulation of XPG/XPF-mediated DSBs at satellite and rDNA repeats in ICF cells

Since the loss-of-function mutation of DNMT3b is the cause of type 1 ICF, we then tested whether R-loops also cause DNA damages in ICF cells. To this end, we first compared endogenous DNA damage by γH2AX IF staining in EBV-immortalized lymphoblastoid cell lines (LCLs) from wild-type and type 1 ICF patient carrying *DNMT3b* mutation (Fig. 4a). The results showed that the levels of γH2AX foci were higher in ICF than those in wild-type LCLs. Similar results were found when comparing fibroblasts from the same ICF patient with normal human IMR-90 fibroblasts (supplementary Fig.S4). The causal role of R-loops in DSBs in ICF cell was tested by expressing RNase H1 using retroviral infection. The amounts of γH2AX foci were clearly decreased by wild-type but not catalytic-dead RNase H1 in ICF cells (Fig.4b). Consistent with γH2AX-ChIP data obtained from BKO cells, expression of wild-type RNase H1 significantly reduced γH2AX association with centromeric, rDNA, and telomeric sequences in these ICF cells, while catalytic-dead RNase H1 had little effect (Fig. 4c). Intergenic region downstream of *SNRPN* that has been shown as an R-loop-free locus (13,30). WT and ICF LCL showed little difference in γH2AX-ChIP at intergenic region downstream of *SNRPN*. These data indicate that R-loops are the source of DNA lesions in ICF LCLs. Thus, the removal of RNA in DNA-RNA hybrid by RNase H1 reduces DNA breaks.

**Figure 4.**
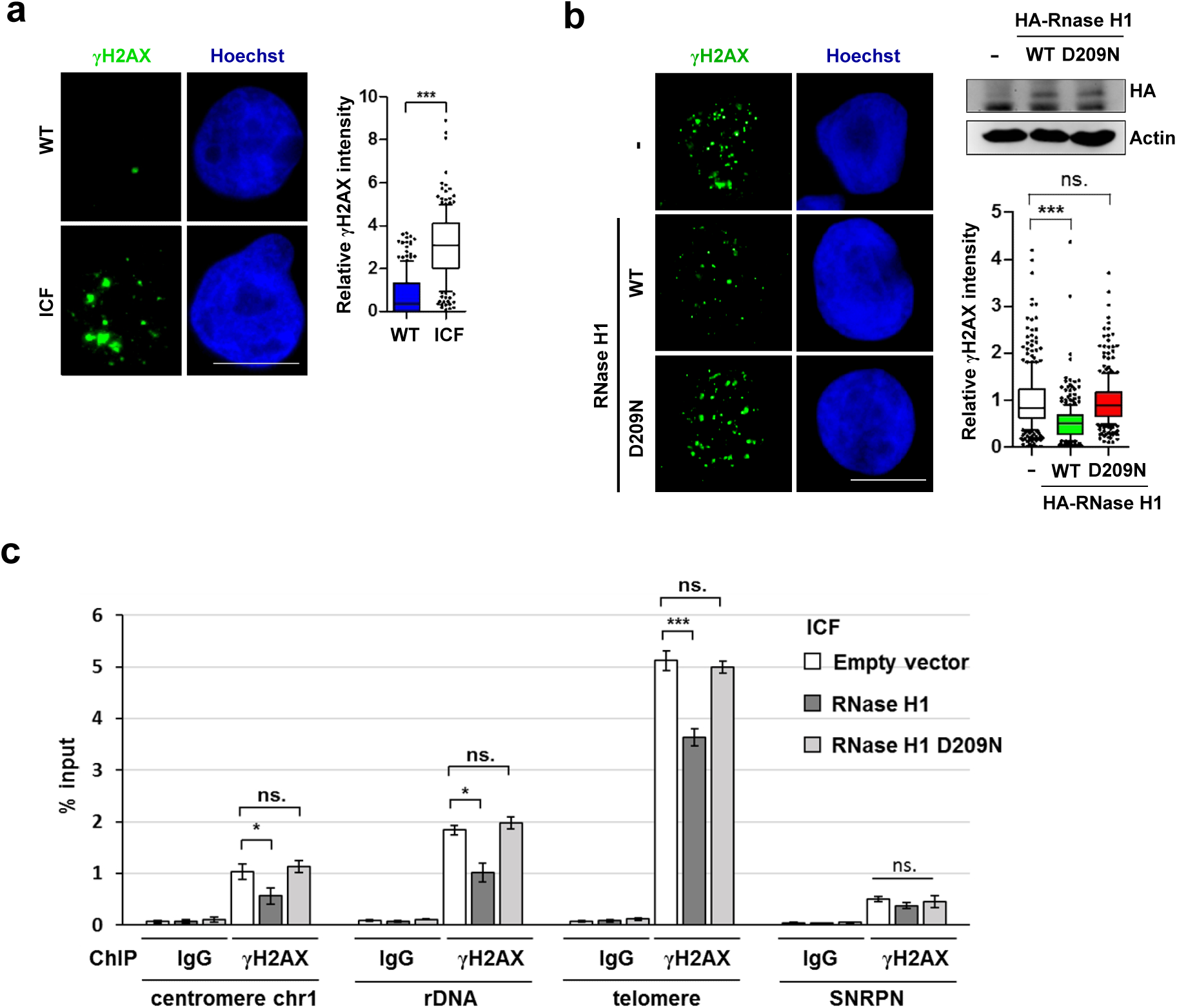
R-loop-mediated DNA breaks at centromeric, rDNA and telomeric sequences in ICF cells. (a) The comparison of DNA damage signal in wild-type vs ICF LCL. Cells were fixed for IF staining by antibody of γH2AX (scale bar, 10μm). Fluorescent intensity of γH2AX in cells (N=100) from three independent experiments was quantitated by Image J and relative intensity is expressed, *\*\*\**P<0.001 by Mann-Whitney test. (b-c) ICF LCLs were infected with retrovirus of empty vector, HA-RNaseH1-WT, or HA-RNaseH1-D209N. (b) γH2AX IF staining, scale bar, 10 μm. Relative fluorescent intensity of γH2AX in cells (N=100) from 3 independent experiments is presented as described in (a). (c) γH2AX-ChIP-qPCR at the sequences of centromere of chromosome1, rDNA, telomere of chromosome 10, and intergenic region downstream of *SNRPN*. Results are expressed as percentage of input DNA in γH2AX versus IgG control. Data are represented as mean ± SEM of three independent experiments and analyzed by two-tailed unpaired Student’s t-test, *, ***P< 0.01, 0.001, ns: no significant. Western blot analysis of HA-RNaseH1-WT and HA-RNaseH1-D209N was shown in the right panel.

Like in BKO cells, the effect of XPG or XPF single knockdown on DSBs in ICF cells was very little (supplementary Fig.S5). Simultaneous knockdown of XPG and XPF was able to abolish γH2AX foci in ICF LCLs (Fig. 5a). Similar results were found in ICF fibroblasts (Supplementary Fig. S6). The comet assay showed the reduction in tail moment by double XPG/XPF knockdown, confirming XPG/XPF-mediated genome damage (Fig. 5b). The γH2AX-ChIP data also revealed that XPG/XPF knockdown reduced γH2AX binding at sequences of centromere of chromosome 1, rDNA, and telomere in ICF (Fig. 5c). Of note, XPG/XPF double knockdown did not affect the protein levels of H2AX, excluding the possibility of their expression regulation by XPG/XPF (Fig. 1e and 5a). Thus, the loss-of-function mutation of *DNMT3b* in ICF cells also promotes R-loop-mediated DNA damage via XPG/XPF cleavage in repetitive satellite and rDNA sequences. As for wild-type LCL cells, the levels of γH2AX associated with sequences of centromere of chromosome 1 and rDNA were low, and knockdown of XPG/XPF had no effect. However, γH2AX signals in telomere was increased by XPG/XPF knockdown. This indicates that XPG/XPF is probably critical for preventing DNA damage at telomere sites in normal LCL cells. This result is probably relevant to a previous report describing that NER is important for maintenance of telomere stability in normal somatic cells (31,32).

**Figure 5.**
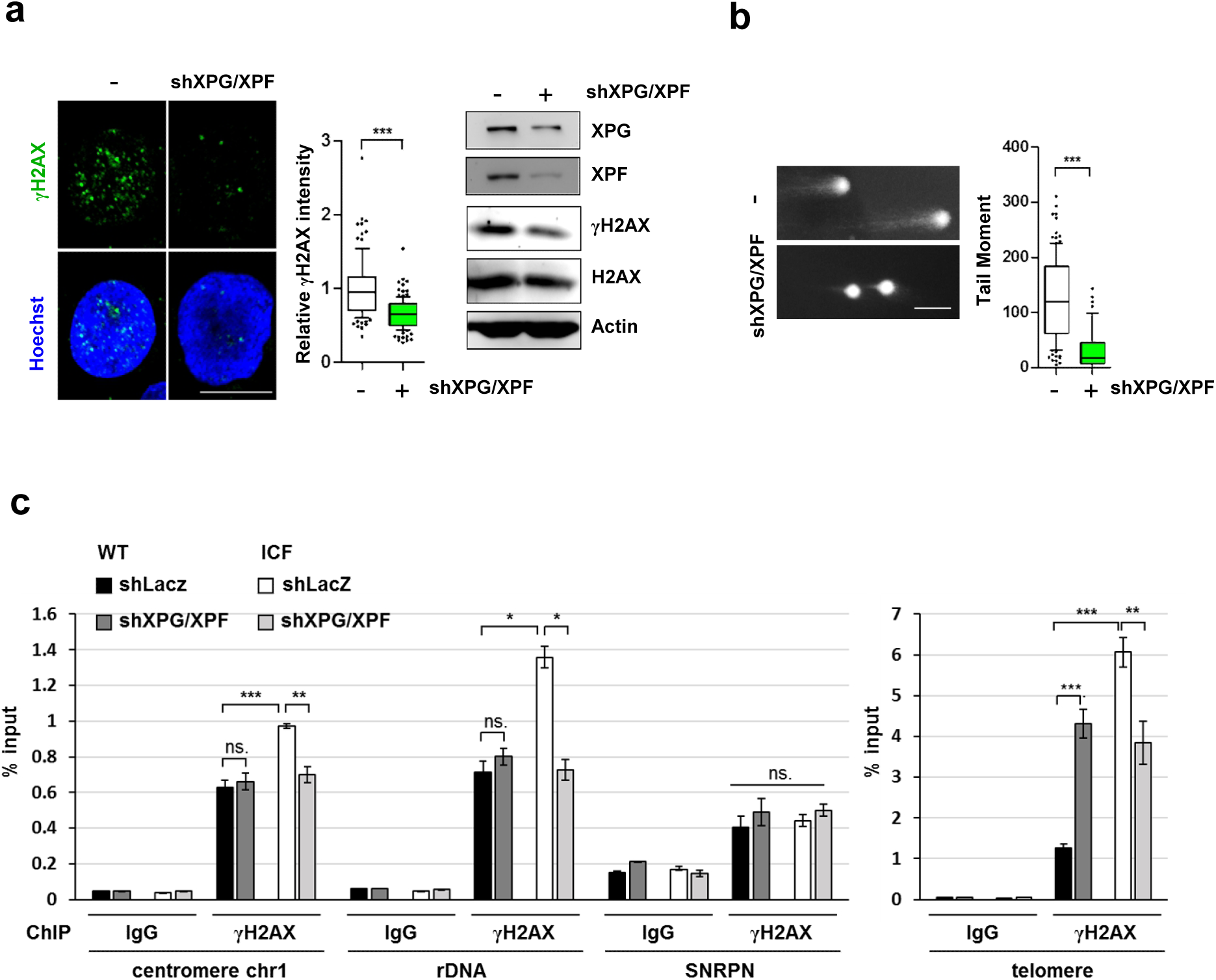
XPG/XPF knockdown reduces DNA breaks in ICF cells. (a-b) ICF cells with or without XPG/XPF knockdown for (a) γH2AX IF staining with Hoechst (scale bar, 10 μm) and western blot analyses, and (b) comet tail analysis. Fluorescent intensity of γH2AX IF staining (N>100) from three independent experiments was quantitated by Image J and relative intensity is expressed, *\*\*\**P<0.001 by Mann-Whitney test. Western blot analysis of XPG, XPF, γH2AX, H2AX, and actin was shown in the right panel. (b) Comet tail moments in cells (N=300) were measured and analyzed by CometScore. (c) γH2AX-ChIP-qPCR in wild-type and ICF LCL cells with or without XPG/XPF knockdown. Data are expressed as described in figure 4c (mean ± SEM, n=3, *, **, *** P < 0.05, 0.01, 0.001, ns: no significant by two-tailed unpaired Student’s t-test).

### Less R-loop and more DNA break at centromeric sites in ICF cells

We further accessed the level of centromeric R-loops in wild-type and ICF LCL cells in response to XPG/XPF knockdown. The DRIP analysis showed the amount of centromeric R-loop detected in ICF cells was much less than that in wild-type cells (Fig. 6a). However, in XPG/XPF knockdown cells, the amount of DNA-RNA hybrid at centromeric site were similar in wild-type and ICF LCLs (Fig. 6a). Very likely, the loss of DNMT3b function in ICF cells causes centromeric R-loops more vulnerable to XPG/XPF-mediated DNA cleavage, thereby making more DNA breaks and less R-loops in centromere. We also used the samples from ICF cells expressing wild-type and catalytic-dead RNase H1 to assure the reliability of our DRIP assays. In accordance with the dominant-negative property of catalytic-dead mutant, expression of RNaseH1 (D209N) elevated centromeric R-loop, validating the R-loop analysis in these cells (supplementary Fig.S7). We also tested whether XPG/XPF knockdown can affect chromosome segregation in ICF and wild-type cells. To this end, cells were treated with nocodazole overnight. By washing out nocodazole to allow mitotic progression, the amounts of lagging and bridge chromosomes in mitosis were evaluated. As compared to wild-type cells, ICF LCL cells had an increased population containing lagging and bridge chromosomes, which were mitigated by XPG/XPF knockdown (Fig. 6b)

**Figure 6.**
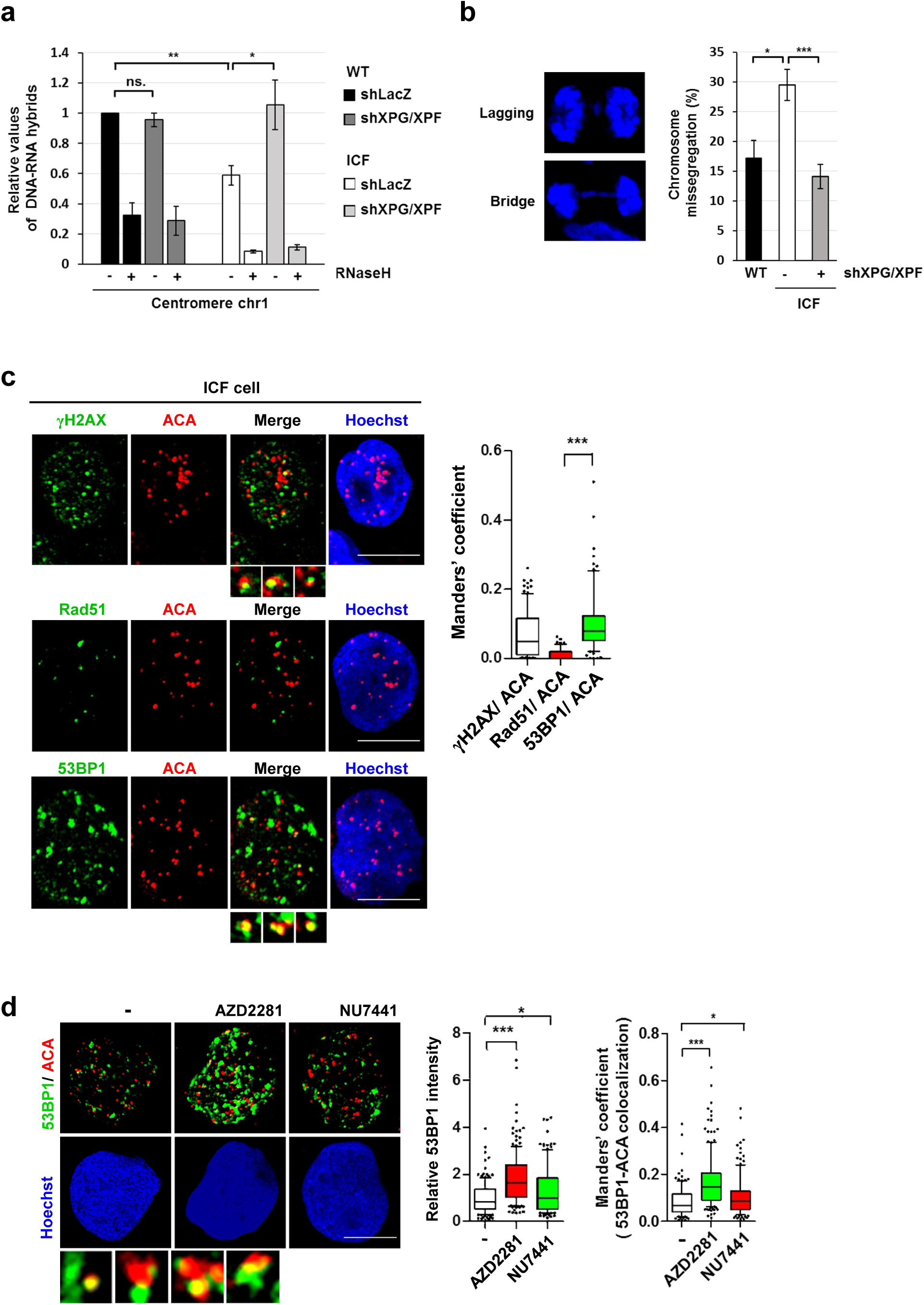
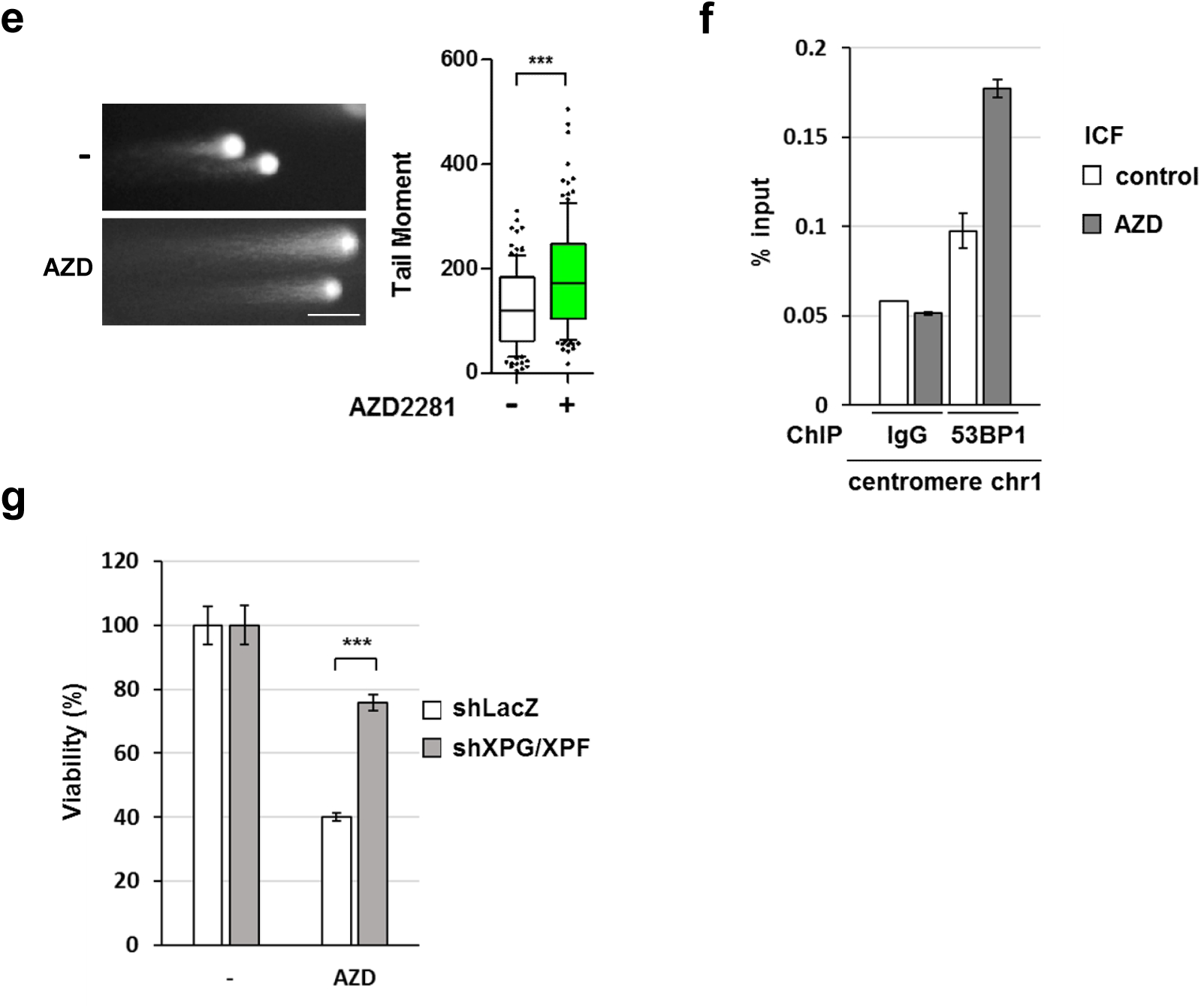
The increased removal of centromeric R-loop and PARP1-mediated NHEJ repair in ICF cells. (a) DRIP analysis of centromeric R-loops in wild-type and ICF LCLs with or without XPG/XPF knockdown. DNA samples were untreated (-) or treated (+) with RNaseH before immunoprecipitated by S.9.6 antibody. qPCR values of DNA-RNA hybrids at centromere sequence of chromosome 1 are normalized by the value at the negative control region of *SNRPN*. Data are expressed relative to WT cells (mean ± SEM, n=3, *, **P< 0.05, 0.01, ns: no significant by two-tailed unpaired Student’s t-test). (b) Wild-type, ICF, and XPG/XPF-depleted ICF LCLs were arrested in mitosis by nocodazole treatment overnight. Cells were released from nocodazole arrest for 70 min to analyze chromosome segregation. Representative images of chromosome lagging and bridge errors are shown. Percentages of cells with anaphase bridges and lagging chromosomes are shown in the histogram. Error bars are shown in means ± SEM, n=3. *, *** indicate P< 0.05 and 0.001, respectively by two-tailed unpaired Student’s t-test. (c) ICF cells were fixed for γH2AX/ACA, Rad51/ACA, and 53BP1/ACA IF co-staining. Representative confocal colocalized signals are shown under the merged image, Hoechst indicates nuclei, scale bar, 10 μm. Manders’ coefficient of colocalized γH2AX/ACA, Rad51/ACA, and 53BP1/ACA was acquired by confocal imaging as described in methods. Examples of co-localized signals of γH2AX/ACA and 53BP1/ACA are showed in the image underneath. Quantitation data of colocalized signals are expressed (N>100), *\*\*\** P<0.001 by Mann-Whitney test. (d) ICF LCLs were treated with NU7441 (5 μM) or AZD2281 (5 μM) for 6h and fixed for 53BP1/ACA IF co-staining, scale bar, 10 μm. The fluorescent intensity of 53BP1 was quantitated by Image J and relative intensity is expressed (N>100), *, *\*\*\**P<0.001 by the Mann-Whitney test. Manders’ coefficient of co-localized 53BP1/ACA was quantified as described in (c). Examples of co-localized signals of 53BP1/ACA are shown in the image underneath. Quantitation data of co-localized signal are expressed (N>150), *, *\*\*\**P <0.05, 0.001 by Mann-Whitney test. (e and f) ICF LCLs were treated with AZD2281 (5 μM) for 6 h. (e) Comet tail moments analysis. Cells (N=300) were measured and analyzed by CometScore (*\*\*\**P<0.001 by Mann-Whitney test). (f) 53BP1-ChIP-qPCR. Data are shown as percentage of input DNA at centromere sequence of chromosome 1 in 53BP1 antibodies versus IgG control in two independent experiments. (g) Viability assay. ICF LCLs with or without shXPG/XPF knockdown were treated with AZD2281 (5 μM) for 72h. The percentage of the viable cell from three independent experiments is expressed, mean ± SEM, ***P<0.001 by two-tailed unpaired Student’s t-test.

### Mutagenic DNA repair in centromere

It has been reported that DNMT3b mutation in ICF leads to centromere shortening (9,10). Presumably, endogenous DSBs are constantly repaired to prevent genome deterioration for cell survival. We suspected that the repair of DNA breaks in centromere leads to shortening. DSBs are repaired by homologous recombination (HR) or non-homologous end-join (NHEJ) repair pathway (33). HRR pathway involves an extended BRCA1-dependent end resection and loading of Rad51 for strand invasion to give error-free repair (34). 53BP1 recruitment to DSB sites antagonizes the HRR pathway and favors NHEJ repair (34-36). By confocal microscopic analysis, γH2AX and 53BP1 were well colocalized with IF signal by anti-centromere antibody (ACA) that marks centromere sites. In contrast, little Rad51 foci were colocalized with ACA signal in ICF cells (Fig.6c), suggesting the lack of ongoing HRR at centromeric sites. We then tested NHEJ in the repair of DNA damage in ICF cells. Two NHEJ pathways, conventional NHEJ (c-NHEJ) and alternative-end joining (Alt-EJ), participate in the repair of two-ended and one-ended DSBs (37). NU7441, an inhibitor of DNA-PK, specifically suppresses c-NHEJ. AZD2281, a PARP1 inhibitor, suppresses alt-EJ and c-NHEJ. ICF cells were then treated with NU7441 and AZD2281 to distinguish the contribution of these two NHEJ pathways in repairing endogenous DSBs. AZD2881 treatment significantly increased overall 53BP1 foci. The amounts of these foci co-localized with ACA were also increased. NU7441 treatment gave a much lesser effect on 53BP1 foci and their colocalization with ACA (Figures 6d). Comet assay also showed that AZD treatment increased the length of the tail moment in ICF cells, confirming the contribution of PARP1 activity in reducing DNA damage in ICF cells (Figure 6e). We further performed ChIP analysis by 53BP1 antibody in ICF LCLs with and without AZD treatment. The result showed that inhibition of PARP1 significantly increased 53BP1 binding at centromere site of chromosome 1 (Figure 6f), suggesting that the centromere DNA breaks are constantly repaired in a PARP1-dependent manner. As such, AZD treatment decreased the viability of ICF cells, whereas XPG/XPF knockdown ICF cells were less susceptible to this treatment (Figure 6g). Likely, inhibition of PARP1 accumulates DSBs in centromere, which become detrimental to the viability of ICF cells. Thus, DSBs at centromeric sites are frequently repaired by PARP-1-mediated NHEJ pathway. Since NHEJ is a mutagenic repair, our data imply that XPG/XPF-induced DSBs in centromere and the participation of NHEJ repair lead to centromere instability in ICF cells.

## DISCUSSION

This study revealed that R-loops-mediated DNA damage is the major cause of pronounced DNA damage signals observed in DNMT3b deficient HCT116 and type 1 ICF cells. It has been found that DNMT3b dysfunction mainly causes hypomethylation in repetitive sequences rather than promoters of coding genes (1,38). Consistently, we found that R-loop-mediated DNA damages are spreading over these repetitive sequences at centromere, telomere, and rDNA regions. Since XPG/XPF knockdown was able to abolish DNA damage signal in these cells, it is apparent that R-loop-mediated DNA breaks in these sequences are the results of DNA cleavages by these two DNA endonucleases in NER process. Importantly, we found that knockdown of XPG or XPF alone was unable to abolish DSBs in either BKO or ICF cells, indicating that the action of either one of XPG- or XPF-mediated cleavage is sufficient to cause DSBs formation. Presumably, each endonuclease acts to generate a nick at one strand of DNA, which would not directly give DSB. Therefore, it is possible that DNA nicks from either XPG or XPF cleavage in each strand within these repetitive sequences collide with replication fork, resulting in one-ended DSBs.

DNA hypomethylation not only affects transcription but also histone modification and recruitment of chromatin factors (39). Our findings further addressed the question of whether the induction of R-loop-mediated DNA damages by DNMT3b dysfunction is through generating too much R-loops or having more DNA cleavages by XPG/XPF. Of note, the steady-state level of centromeric R-loops in ICF is indeed less than that in wild-type LCL. After XPG/XPF knockdown, the level of centromeric R-loops becomes similar in wild-type and ICF LCL. Therefore, the loss-of-function DNMT3b mutation increases the removal of centromeric R-loop by NER process, rather than having excessive amounts of R-loop formation. Accordingly, we proposed that DNA methylation by DNMT3b at centromere R-loop sites mediates an as-yet-unknown process in restricting the recruitment of NER factors, thus preventing R-loops in centromere from XPG/XPF-mediated processing. ICF cells defective in DNMT3b function, therefore, have more DSBs in centromere but less amount of R-loops. In fact, we found that XPG/XPF knockdown did increase ATR binding in centromeric R-loop and mitigate the error of chromosome segregation in BKO cells. Thus, XPG/XPF-mediated processing of centromeric R-loops acts as a double-edged sword in chromosome instability by eliminating mitotic checkpoint function and increasing DNA breaks.

This study also investigated the repair of these R-loops-mediated DSBs in centromere. A previous study has shown that the HR repair factor is excluded from the heterochromatin domain to prevent abnormal recombination of repetitive sequences unless the DSB site moves outside the domain (40). Although conventional NHEJ repair can cause small deletion to threaten the integrity of the coding gene, this problem might not be severe enough to affect repetitive satellite DNA sequences that span over the mega-base range. PARP1 is critical for c-NHEJ and alt-EJ pathways, while DNA-PK is required for c-NHEJ but not alt-EJ repair (41-43). In this study, inhibition of DNA-PK had a rather moderate effect on the overall 53BP1 foci, whereas PARP1 inhibition markedly increased these DSB signals. A recent study has demonstrated that c-NHEJ function is compromised in type 3 and type 4 ICF associated with mutation of CDCA7 and HELLS. Therefore, these patient cells have increased level of γH2AX and centromere instability. In the report, they also showed that the cell viability of two DNMT3b knockout clones of HEK293 cells are not affected by DNA-PK inhibitor treatment, while the cell viability of one clone was affected by PARP1 inhibitor (44). In this study, alt-EJ pathway rather than c-NHEJ is more involved in the repair of XPG/XPF-mediated DNA damage at the repetitive centromere sequences in type 1 ICF cell. Our ChIP analysis further showed that 53BP1 binding at centromeric sites was increased by PARP1 inhibition. Thus, PARP1-dependent repair of XPG/XPF-mediated DNA breaks plays an important role in eliminating the frequent breakages at centromere for cell survival in ICF cells. Accordingly, we hypothesize that centromeric DNA breaks generated by XPG/XPF-mediated removal of R-loops are repaired by NHEJ to cause the end-joining at break sites, leading to centromere shortening, metaphase quadriradials with mixed chromosomal arms as common features observed in ICF due to DNMT3b dysfunction (9,10).

## MATERIAL AND METHODS

### Cell culture, transfection, and infection

HCT116 and BKO cells were kindly provided by Bert Vogelstein (Johns Hopkins University School of Medicine, MD, USA) and were maintained in McCoy’s 5A medium supplemented with 10% fetal bovine serum (FBS), 100 U/ml penicillin and 10 mg/ml streptomycin. Type-1 ICF cells, pGM08747 (fibroblasts), and pGM08714 (lymphoblastoid cell lines, LCLs) and normal human fibroblasts, IMR-90, were obtained from Coriell Cell Repositories. Normal (WT) LCLs were kindly provided by Prof. Ching-Hwa Tsai (Graduate Institute of Microbiology, National Taiwan University). LCLs were maintained in RPMI supplemented with 15% FBS, 2 mM glutamine, 100 U/ml penicillin, and 10 mg/ml streptomycin. IMR-90 and ICF fibroblasts were grown in Dulbecco’s modified Eagle’s medium supplemented with 15% FBS, 100 U/ml penicillin, and 10 mg/ml streptomycin. HEK293T and Platinum-A retroviral packaging (Cell Biolabs, CA, USA) cells were maintained in Dulbecco’s modified Eagle’s medium supplemented with 10% FBS, 100 U/ml penicillin and 10 mg/ml streptomycin. HEK293T cells were cotransfected with pCMVdeltaR8.91, pCMV VSVG, and targeted plasmids including PL-SIN-5TO-p53-IRES-GFP, *ERCC4* (XPF) shRNA (TRCN0000078587) and *ERCC5* (XPG) shRNA (TRCN0000358878) to produce lentivirus. For LCL cells, individual shRNA viral infection was performed by incubation of cells with viral supernatant followed by centrifugation at 800 x g for 1 hour and recovery for 2 days. For double knockdown by shXPG and shXPF viral infection, cells were infected with shXPG virus and selected with puromycin (1 mg/ml), followed by subsequent shXPF viral infection. After recovery for 2 days, cells were incubated in the growth medium containing puromycin (1 mg/ml). For retrovectors infection, pMXs-3XHA-mouseRNaseH1 and pMXs-3XHA-mouseRNaseH1-D209N, were transfected into Platinum-A cells to produce retrovirus. After transfection for 48h, the supernatants containing virus were collected for infection.

### Antibodies and reagents

Antibodies: anti-DNA-RNA hybrid S9.6 (Millipore, MABE1095), HA (Santa Cruz, sc-805), β-tubulin (Sigma-Aldrich, T4026), β-Actin (Santa Cruz, sc-8432), Chk1 (Santa Cruz, sc-8408), phospho S345-Chk1 (cell signaling, #2341), Chk2 (Millipore, 05-649), phospho T68-Chk2 (cell signaling, #2661), phospho S1981-ATM (Gene Tex, GTX61739), ATR (Bethyl Laboratories, A300-128A), phospho S139-H2AX (Millipore, 05-636), phospho S139-H2AX (Abcam, ab2893), H2AX (Gene Tex, GTX127340), H3S10p (Millipore, 06-570), XPF (Santa Cruz, sc-398032), XPG (Proteintech, 11331-1-AP), CSB (Santa Cruz, sc-166042), and 53BP1 (Millipore, MAB3802). Chemicals for cell treatment: Nocodazole (Sigma, M1404), NU7741 (Santa Cruz, sc-208107), and AZD2281 (Selleckchem, S1060).

### Immunofluorescence staining

Cells were fixed with 4% paraformaldehyde for 15 min at room temperature (RT), followed by blocking in 5% BSA/TBS for 1h at RT and staining with primary antibody overnight at 4°C. After incubation with secondary antibody for 1 hr at RT, slides were mounted in Fluoro-gel mounting oil (EMS, #17985-10) and were analyzed by an OLYMPUS BX53 and LSM 700 laser scanning confocal microscope (Carl Zeiss).

### Comet assay

Comet assay was performed by Electrophoresis Assay (Trevigen, Inc) according to the manufacturer’s protocol. DNA was stained with ethidium bromide and analyzed with Image J (v 1.47) for measuring tail length (TL). The comet tail (TM = %DNA in tail × TL/100) according to the manufacturer’s suggestion.

### Chromatin immunoprecipitation (ChIP)-sequencing

Chromatin immunoprecipitation using the anti-gamma H2A.X (Abcam, ab2893) was carried out according to a previously described protocol (45). Libraries were constructed and bar-coded using TruSeq RS Cluster kit-HS (Illumina) and single-end sequencing (50 bp) was performed using an Illumina HiSeq-2500 sequencer at Genomic Research Center of National Yang-Ming University (Taiwan) according to manufacturer’s instruction. All reads were mapped to the human genome (hg38) using the bowtie2 alignment software (46). The alignment results were used to call peaks by MACS (47). The results of the peak signal were subject to the Integrative Genomics Viewer (IGV, Broad Institute) (48) for further comparison and analysis. We used UCSC hg38 and Rfam v11 to identify and annotate the ChIP-seq peak regions.

### DNA-RNA immunoprecipitation (DRIP) assay

DRIP was performed as described in (49). Briefly, DNA was extracted carefully by Wizard® Genomic DNA Purification Kit (Promega, A1125), washed with 70% EtOH, and resuspended in TE buffer. The purified nucleic acids were sonicated in a buffer containing 10 mM Tris-HCl pH 8.5 and 300 mM NaCl to yield an average DNA fragment size of ∼300 bp. Sonicated DNA (6μg) was added in the IP buffer (50 mM Hepes/KOH at pH 7.5; 0.14 M NaCl; 5 mM EDTA; 1% Triton X-100; 0.1% Na-Deoxycholate), followed by addition of S9.6 antibody (10μg) for incubation overnight at 4°C. For removal of R-loops as a negative control, 6μg of sonicated DNA was pretreated with 12 μl of RNase H (5000 U/mL; NEB) for incubation at 37°C for 6h prior to immunoprecipitation by S9.6 antibody. Protein G Sepharose beads that were pre-blocked with PBS containing 0.5% BSA, were added for 4h to pull down immunocomplex. Bound beads were recovered and washed with 1ml of low salt buffer (50mM Hepes/KOH, pH 7.5, 0.14 M NaCl, 5 mM EDTA pH 8, 1% Triton X-100, 0.1% Na-Deoxycholate), 1 ml of lysis buffer (50 mM Hepes/KOH pH 7.5, 0.5 M NaCl, 5 mM EDTA pH 8, 1% Triton X-100, 0.1% Na-Deoxycholate), 1 ml of washing buffer (10 mM Tris-HCl, pH 8, 0.25 M LiCl, 0.5% NP-40, 0.5% Na-Deoxycholate, 1 mM EDTA pH 8), and 1 ml of TE (100 mM Tris-HCl, pH 8, 10 mM EDTA, pH 8) at 4°C. Precipitates were eluted in elution buffer (10 mM Tris pH 8, 1 mM EDTA, 1% SDS) in 100 μl for 15 min at 65°C. DNA was purified with QIAquick® PCR purification Kit (QIAGEN, 28106).

### qPCR for ChIP and DRIP

qPCR was performed on a StepOne™ Real-Time PCR System (Applied Biosystems™ LS4376357) using the Fast SYBR-Green master mix (Applied Biosystems™). qPCR primers are as followed:

Chr1 centromere, forward TCATTCCCACAAACTGCGTTG and reverse TCCAACGAAGGCCACAAGA; Chr4 centromere forward GTGGGAACCACA GAACCACT and reverse TTTCATGCGCCACCTTTTGG; Chr10 telomere, forward GTCCGTCCGTGAAATTGCG and reverse GGTCCAAACGAGTCTCCGTC; rDNA, forward CGATGGTGGCGTTTTTGG and reverse CCGACTCGGAGCGAAAGATA; p21, forward CTGCCCAAGCTCTACCTTCCCA and reverse GGTCCACATGGTCTTCCTCTGC; *SNRPN* as a control, forward GCCAAATGAGTGAGGATGGT and reverse TCCTCTCTGCCTGACTCC AT.

### DNA fiber analysis

DNA fiber assay was performed as described previously (50). Briefly, cells were incubated medium containing 25 mM of CldU (Sigma-Aldrich, C6891) for 30 min, followed by replacement with a medium containing 250 mM of IdU (Sigma-Aldrich, I7125) for 30 min. Cells were harvested and suspended in a solution (200 mM Tris-HCl, 50 mM EDTA, and 0.5% SDS). After spreading on coverslip and air-dried, the coverslip was fixed with methanol/acetic acid (3:1), denaturation, and blocking, prior to staining with rat anti-BrdU antibody (which detects CldU but not IdU, OBT0030, 1:2,000, AbDSerotec) and mouse anti-BrdU antibody (which detects IdU but not CldU, 7580, 1:1,000, BD Biosciences). Images of DNA fibers were acquired using a microscope (OLYMPUS BX53). The lengths of red- and green-labeled fibers were determined by FluoView3.0 software (Olympus).

### Chromosome segregation analysis

Cells were treated with 0.2 μg/ml nocodazole for 16h. After replacing with fresh medium for 1h, cells were fixed with 4% paraformaldehyde for 15 min at RT, followed by blocking in 5% BSA/TBS for 1h at RT and staining with mouse antibody to β-tubulin (1:200) and rabbit antibody to phospho Ser10-H3 (1:500) overnight at 4°C. Cells were incubated with secondary antibodies in the presence of Hoechst 33342 for 1 hour at RT. Cells were mounted in Fluoro-gel mounting oil (EMS, #17985-10) for observation by OLYMPUS BX53 and confocal microscope (ZEISS, LSM700).

### Confocal microscope and image quantification

Co-localization of two proteins of IF staining was acquired by LSM 700 laser scanning confocal microscope (Carl Zeiss). The conditions of IF staining, laser power, and pinhole sizes by confocal microscopy were identical among groups. Pixel number, pixel intensity, and area were provided by built-in software in LSM 700. Manders’ coefficient was calculated for the co-localization of two proteins in the nuclear area. The co-localization coefficient values are ranged from 0 (segregation) to 1 (complete co-localization).

### Cell viability assay

Cells were plated at 1,000 cells per well in a 96-well plate. Cells were treated with NU7441 (5μM) or AZD2281 (5μM) for 72 h, after which 10 μl of the Cell Counting Kit-8 (CCK-8) solution was added into each well for an additional 2 h at 37°C. The absorbance at 450 nm was measured using a Tecan Spark multimode microplate reader and the percentage of cell viability was calculated.

### Statistics

Statistical analysis was performed using Student’s t-test and Mann-Whitney test. Differences were considered statistically significant when P values< 0.05, 0.01, and 0.001 indicated by *, **, and ***, respectively. Error bars represent the SEM of at least three independent experiments.

### Data availability

Sequencing data have been deposited at the GEO under the accession number GSE142376 and linked to UCSC genome browser session of DNMT3B study. https://genome.ucsc.edu/s/chenwy/rH2AX_ChIP%2Dseq_HCT116

## ACKNOWLEDGMENTS AND FUNDINGS

We thank Prof. Ching-Hwa Tsai for providing normal (WT) immortalized LCLs. This study was financially supported by the “Center of Precision Medicine” from The Featured Areas Research Center Program within the framework of the Higher Education Sprout Project by the Ministry of Education (MOE) in Taiwan and by the grant, MOST 107-3017-F-002-002 and MOST 107-2321-B-002-041 from the Ministry of Science and Technology, Taiwan, (R.O.C.).

## AUTHOR CONTRIBUTIONS

HTS designed, performed experiments, and analyzed data. WYC did ChIP-seq, RNA-seq experiments, and HA-RNase H1 cloning. HYW did chromosome experiments. HDH and CHC performed all bioinformatics analyses. ZFC wrote the manuscript. ZFC supervised the entire project.

## Notes

### Competing Interest Statement

The authors have declared no competing interest.

## REFERENCES

1. Okano, M., Bell, D. W., Haber, D. A., and Li, E. (1999) DNA methyltransferases Dnmt3a and Dnmt3b are essential for de novo methylation and mammalian development. Cell 99, 247–257

2. Li, E., Bestor, T. H., and Jaenisch, R. (1992) Targeted mutation of the DNA methyltransferase gene results in embryonic lethality. Cell 69, 915–926

3. Dodge, J. E., Okano, M., Dick, F., Tsujimoto, N., Chen, T., Wang, S., Ueda, Y., Dyson, N., and Li, E. (2005) Inactivation of Dnmt3b in mouse embryonic fibroblasts results in DNA hypomethylation, chromosomal instability, and spontaneous immortalization. J Biol Chem 280, 17986–17991

4. Xu, G. L., Bestor, T. H., Bourc’his, D., Hsieh, C. L., Tommerup, N., Bugge, M., Hulten, M., Qu, X., Russo, J. J., and Viegas-Pequignot, E. (1999) Chromosome instability and immunodeficiency syndrome caused by mutations in a DNA methyltransferase gene. Nature 402, 187–191

5. Hansen, R. S., Wijmenga, C., Luo, P., Stanek, A. M., Canfield, T. K., Weemaes, C. M., and Gartler, S. M. (1999) The DNMT3B DNA methyltransferase gene is mutated in the ICF immunodeficiency syndrome. Proc Natl Acad Sci U S A 96, 14412–14417

6. Gisselsson, D., Shao, C., Tuck-Muller, C. M., Sogorovic, S., Palsson, E., Smeets, D., and Ehrlich, M. (2005) Interphase chromosomal abnormalities and mitotic missegregation of hypomethylated sequences in ICF syndrome cells. Chromosoma 114, 118–126

7. Gopalakrishnan, S., Sullivan, B. A., Trazzi, S., Della Valle, G., and Robertson, K. D. (2009) DNMT3B interacts with constitutive centromere protein CENP-C to modulate DNA methylation and the histone code at centromeric regions. Hum Mol Genet 18, 3178–3193

8. Heyn, H., Vidal, E., Sayols, S., Sanchez-Mut, J. V., Moran, S., Medina, I., Sandoval, J., Simo-Riudalbas, L., Szczesna, K., Huertas, D., Gatto, S., Matarazzo, M. R., Dopazo, J., and Esteller, M. (2012) Whole-genome bisulfite DNA sequencing of a DNMT3B mutant patient. Epigenetics 7, 542–550

9. Jaco, I., Canela, A., Vera, E., and Blasco, M. A. (2008) Centromere mitotic recombination in mammalian cells. J Cell Biol 181, 885–892

10. Barra, V., and Fachinetti, D. (2018) The dark side of centromeres: types, causes and consequences of structural abnormalities implicating centromeric DNA. Nat Commun 9, 4340

11. Bersani, F., Lee, E., Kharchenko, P. V., Xu, A. W., Liu, M., Xega, K., MacKenzie, O. C., Brannigan, B. W., Wittner, B. S., Jung, H., Ramaswamy, S., Park, P. J., Maheswaran, S., Ting, D. T., and Haber, D. A. (2015) Pericentromeric satellite repeat expansions through RNA-derived DNA intermediates in cancer. Proc Natl Acad Sci U S A 112, 15148–15153

12. Chan, F. L., Marshall, O. J., Saffery, R., Kim, B. W., Earle, E., Choo, K. H., and Wong, L. H. (2012) Active transcription and essential role of RNA polymerase II at the centromere during mitosis. Proc Natl Acad Sci U S A 109, 1979–1984

13. Kabeche, L., Nguyen, H. D., Buisson, R., and Zou, L. (2018) A mitosis-specific and R loop-driven ATR pathway promotes faithful chromosome segregation. Science 359, 108–114

14. Crossley, M. P., Bocek, M., and Cimprich, K. A. (2019) R-Loops as Cellular Regulators and Genomic Threats. Mol Cell 73, 398–411

15. Aguilera, A., and Gomez-Gonzalez, B. (2017) DNA-RNA hybrids: the risks of DNA breakage during transcription. Nat Struct Mol Biol 24, 439–443

16. Santos-Pereira, J. M., and Aguilera, A. (2015) R loops: new modulators of genome dynamics and function. Nat Rev Genet 16, 583–597

17. Bhatia, V., Herrera-Moyano, E., Aguilera, A., and Gomez-Gonzalez, B. (2017) The Role of Replication-Associated Repair Factors on R-Loops. Genes (Basel) 8

18. Bonnet, A., Grosso, A. R., Elkaoutari, A., Coleno, E., Presle, A., Sridhara, S. C., Janbon, G., Geli, V., de Almeida, S. F., and Palancade, B. (2017) Introns Protect Eukaryotic Genomes from Transcription-Associated Genetic Instability. Mol Cell 67, 608–621 e606

19. Sollier, J., Stork, C. T., Garcia-Rubio, M. L., Paulsen, R. D., Aguilera, A., and Cimprich, K. A. (2014) Transcription-coupled nucleotide excision repair factors promote R-loop-induced genome instability. Mol Cell 56, 777–785

20. Cristini, A., Ricci, G., Britton, S., Salimbeni, S., Huang, S. N., Marinello, J., Calsou, P., Pommier, Y., Favre, G., Capranico, G., Gromak, N., and Sordet, O. (2019) Dual Processing of R-Loops and Topoisomerase I Induces Transcription-Dependent DNA Double-Strand Breaks. Cell Rep 28, 3167–3181 e3166

21. Garcia-Pichardo, D., Canas, J. C., Garcia-Rubio, M. L., Gomez-Gonzalez, B., Rondon, A. G., and Aguilera, A. (2017) Histone Mutants Separate R Loop Formation from Genome Instability Induction. Mol Cell 66, 597–609 e595

22. Castellano-Pozo, M., Santos-Pereira, J. M., Rondon, A. G., Barroso, S., Andujar, E., Perez-Alegre, M., Garcia-Muse, T., and Aguilera, A. (2013) R loops are linked to histone H3 S10 phosphorylation and chromatin condensation. Mol Cell 52, 583–590

23. Colak, D., Zaninovic, N., Cohen, M. S., Rosenwaks, Z., Yang, W. Y., Gerhardt, J., Disney, M. D., and Jaffrey, S. R. (2014) Promoter-bound trinucleotide repeat mRNA drives epigenetic silencing in fragile X syndrome. Science 343, 1002–1005

24. Groh, M., Lufino, M. M., Wade-Martins, R., and Gromak, N. (2014) R-loops associated with triplet repeat expansions promote gene silencing in Friedreich ataxia and fragile X syndrome. PLoS Genet 10, e1004318

25. Rhee, I., Bachman, K. E., Park, B. H., Jair, K. W., Yen, R. W., Schuebel, K. E., Cui, H., Feinberg, A. P., Lengauer, C., Kinzler, K. W., Baylin, S. B., and Vogelstein, B. (2002) DNMT1 and DNMT3b cooperate to silence genes in human cancer cells. Nature 416, 552–556

26. Rodgers, K., and McVey, M. (2016) Error-Prone Repair of DNA Double-Strand Breaks. J Cell Physiol 231, 15–24

27. Helmrich, A., Ballarino, M., Nudler, E., and Tora, L. (2013) Transcription-replication encounters, consequences and genomic instability. Nat Struct Mol Biol 20, 412–418

28. Sagie, S., Toubiana, S., Hartono, S. R., Katzir, H., Tzur-Gilat, A., Havazelet, S., Francastel, C., Velasco, G., Chedin, F., and Selig, S. (2017) Telomeres in ICF syndrome cells are vulnerable to DNA damage due to elevated DNA:RNA hybrids. Nat Commun 8, 14015

29. Gagnon-Kugler, T., Langlois, F., Stefanovsky, V., Lessard, F., and Moss, T. (2009) Loss of human ribosomal gene CpG methylation enhances cryptic RNA polymerase II transcription and disrupts ribosomal RNA processing. Mol Cell 35, 414–425

30. Bhatia, V., Barroso, S. I., Garcia-Rubio, M. L., Tumini, E., Herrera-Moyano, E., and Aguilera, A. (2014) BRCA2 prevents R-loop accumulation and associates with TREX-2 mRNA export factor PCID2. Nature 511, 362–365

31. Kruk, P. A., Rampino, N. J., and Bohr, V. A. (1995) DNA damage and repair in telomeres: relation to aging. Proc Natl Acad Sci U S A 92, 258–262

32. Rochette, P. J., and Brash, D. E. (2010) Human telomeres are hypersensitive to UV-induced DNA Damage and refractory to repair. PLoS Genet 6, e1000926

33. Chapman, J. R., Taylor, M. R., and Boulton, S. J. (2012) Playing the end game: DNA double-strand break repair pathway choice. Mol Cell 47, 497–510

34. Bunting, S. F., Callen, E., Wong, N., Chen, H. T., Polato, F., Gunn, A., Bothmer, A., Feldhahn, N., Fernandez-Capetillo, O., Cao, L., Xu, X., Deng, C. X., Finkel, T., Nussenzweig, M., Stark, J. M., and Nussenzweig, A. (2010) 53BP1 inhibits homologous recombination in Brca1-deficient cells by blocking resection of DNA breaks. Cell 141, 243–254

35. Kass, E. M., Moynahan, M. E., and Jasin, M. (2010) Loss of 53BP1 is a gain for BRCA1 mutant cells. Cancer Cell 17, 423–425

36. Chapman, J. R., Sossick, A. J., Boulton, S. J., and Jackson, S. P. (2012) BRCA1-associated exclusion of 53BP1 from DNA damage sites underlies temporal control of DNA repair. J Cell Sci 125, 3529–3534

37. Ceccaldi, R., Rondinelli, B., and D’Andrea, A. D. (2016) Repair Pathway Choices and Consequences at the Double-Strand Break. Trends Cell Biol 26, 52–64

38. Ehrlich, M. (2003) The ICF syndrome, a DNA methyltransferase 3B deficiency and immunodeficiency disease. Clin Immunol 109, 17–28

39. Greenberg, M. V. C., and Bourc’his, D. (2019) The diverse roles of DNA methylation in mammalian development and disease. Nat Rev Mol Cell Biol 20, 590–607

40. Chiolo, I., Minoda, A., Colmenares, S. U., Polyzos, A., Costes, S. V., and Karpen, G. H. (2011) Double-strand breaks in heterochromatin move outside of a dynamic HP1a domain to complete recombinational repair. Cell 144, 732–744

41. Ruscetti, T., Lehnert, B. E., Halbrook, J., Le Trong, H., Hoekstra, M. F., Chen, D. J., and Peterson, S. R. (1998) Stimulation of the DNA-dependent protein kinase by poly(ADP-ribose) polymerase. J Biol Chem 273, 14461–14467

42. Spagnolo, L., Barbeau, J., Curtin, N. J., Morris, E. P., and Pearl, L. H. (2012) Visualization of a DNA-PK/PARP1 complex. Nucleic Acids Res 40, 4168–4177

43. Wei, H., and Yu, X. (2016) Functions of PARylation in DNA Damage Repair Pathways. Genomics Proteomics Bioinformatics 14, 131–139

44. Unoki, M., Funabiki, H., Velasco, G., Francastel, C., and Sasaki, H. (2019) CDCA7 and HELLS mutations undermine nonhomologous end joining in centromeric instability syndrome. J Clin Invest 129, 78–92

45. Lu, F., Chen, H. S., Kossenkov, A. V., DeWispeleare, K., Won, K. J., and Lieberman, P. M. (2016) EBNA2 Drives Formation of New Chromosome Binding Sites and Target Genes for B-Cell Master Regulatory Transcription Factors RBP-jkappa and EBF1. PLoS Pathog 12, e1005339

46. Langmead, B., and Salzberg, S. L. (2012) Fast gapped-read alignment with Bowtie 2. Nat Methods 9, 357–359

47. Zhang, Y., Liu, T., Meyer, C. A., Eeckhoute, J., Johnson, D. S., Bernstein, B. E., Nusbaum, C., Myers, R. M., Brown, M., Li, W., and Liu, X. S. (2008) Model-based analysis of ChIP-Seq (MACS). Genome Biol 9, R137

48. Robinson, J. T., Thorvaldsdottir, H., Winckler, W., Guttman, M., Lander, E. S., Getz, G., and Mesirov, J. P. (2011) Integrative genomics viewer. Nat Biotechnol 29, 24–26

49. Halasz, L., Karanyi, Z., Boros-Olah, B., Kuik-Rozsa, T., Sipos, E., Nagy, E., Mosolygo, L. A., Mazlo, A., Rajnavolgyi, E., Halmos, G., and Szekvolgyi, L. (2017) RNA-DNA hybrid (R-loop) immunoprecipitation mapping: an analytical workflow to evaluate inherent biases. Genome Res 27, 1063–1073

50. Chen, C. W., Tsao, N., Huang, L. Y., Yen, Y., Liu, X., Lehman, C., Wang, Y. H., Tseng, M. C., Chen, Y. J., Ho, Y. C., Chen, C. F., and Chang, Z. F. (2016) The Impact of dUTPase on Ribonucleotide Reductase-Induced Genome Instability in Cancer Cells. Cell Rep 16, 1287–1299

