## Supplemental figures for "DNMT3b Dysfunction Promotes DNA cleavages at Centromeric R-loops to Increase Centromere Instability"

### Supplemental data

#### Figure S1

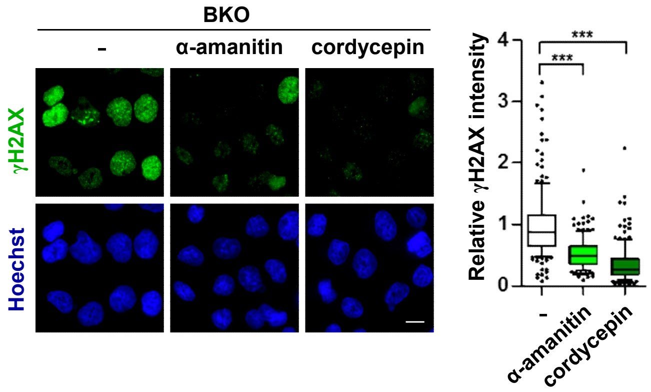


**Supplemental Fig. S1**

BKO cells were treated with RNA polymerase inhibitors α-amanitin (20 μg/ml) or cordycepin (50 μM) for 6h, followed by γH2AX IF staining with Hoechst (scale bar, 10 μm). Fluorescent intensity of γH2AX in cells (N>150) from three independent experiments was quantitated and relative intensity is expressed, ***P<0.001 by Mann-Whitney test.

### Figure S2

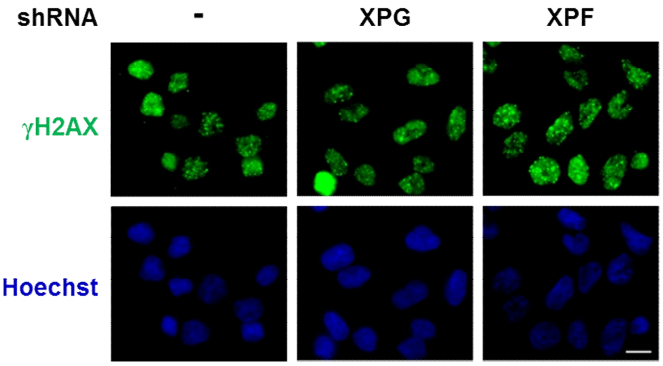


**Supplemental Fig. S2**

BKO cells were infected with shRNA lentivirus of XPG and XPF for γH2AX IF staining. Hoechst indicates nuclei (scale bar, 10 μm).

### Figure S3

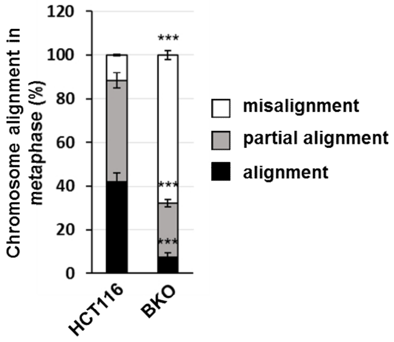


**Supplemental Fig. S3**

Mitotic progression in HCT116 and BKO cells. Cells display aligned, misaligned and partial misaligned metaphase chromatin are defined in Figure 3c. More than 100 cells were counted in each experiment (means ± SEM, three independent experiments. ***P< 0.001 by two-tailed unpaired Student's t-test).

### Figure S4

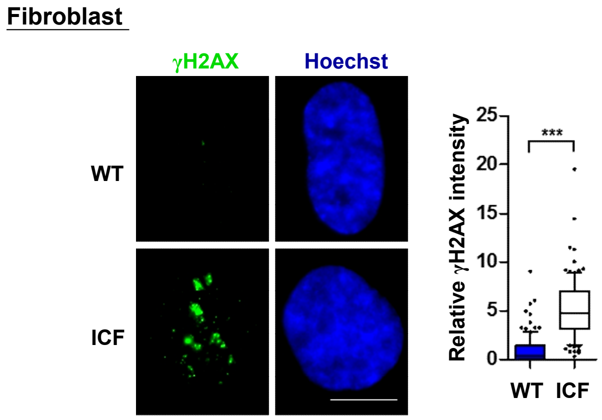


**Supplemental Fig. S4**

Normal IMR-90 and ICF fibroblasts were fixed for IF staining by the antibody of γH2AX. Hoechst indicates nuclei (scale bar, 10 μm). Fluorescent intensity of γH2AX in cells (N>150) from three independent experiments was quantitated by Image J and relative intensity is expressed, ***P<0.001 by the Mann-Whitney test.

### Figure S5

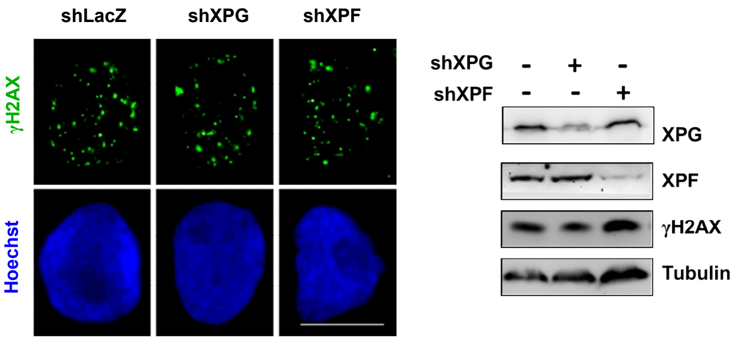


#### Supplemental Fig. S5

ICF LCLs were infected with shRNA lentivirus of XPG and XPF for γH2AX IF staining with Hoechst (scale bar, 10 μm) and western blot analyses. Western blot analysis of XPG, XPF, γH2AX and tubulin is shown in the right panel.

### Figure S6

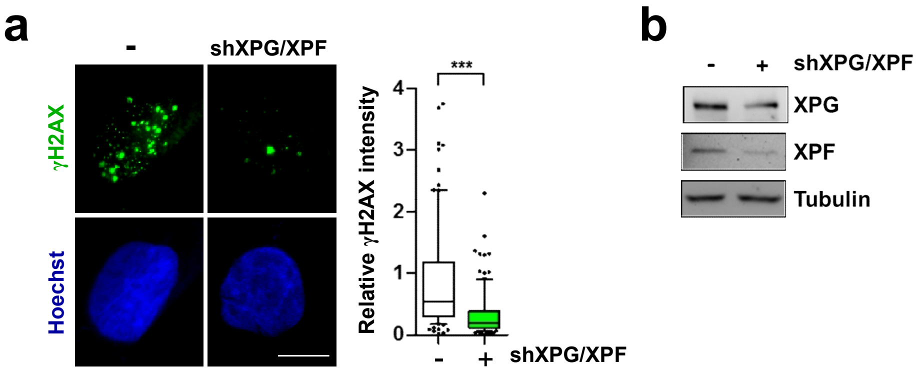


**Supplemental Fig. S6**

(a-b) ICF fibroblast cells with or without XPF/XPG knockdown for (a) γH2AX IF staining with Hoechst (scale bar, 10 μm). Fluorescent intensity of γH2AX in cells (N>50) from three independent experiments was quantitated by Image J and relative intensity is expressed, ***P<0.001 by the Mann-Whitney test. (b) The efficacy for the knockdown of XPG and XPF was shown by western blot.

### Figure S7

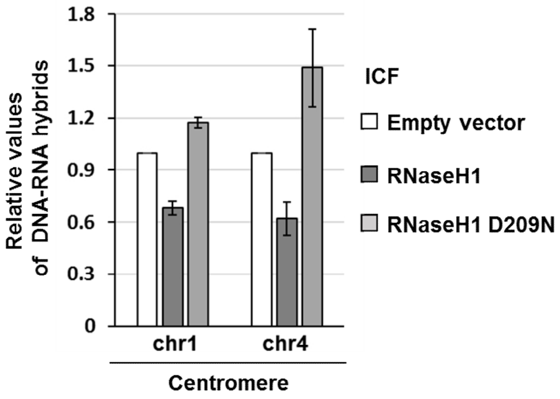


**Supplemental Fig. S7**

ICF LCLs were infected with retrovirus of empty vector, HA-RNaseH1-WT and HA-RNaseH1-D209N for DRIP-qPCR analysis at centromere of chromosome 1 and 4 as indicated. Value of DRIP-qPCR was normalized to the control region of *SNRPN*. Data are expressed relative to empty vector cells from two independent experiments.
